# Integrating Multi-Structure Covalent Docking with Machine Learning Consensus Scoring Enhances Virtual Screening of Human Acetylcholinesterase Inhibitors

**DOI:** 10.1101/2025.09.29.679420

**Authors:** Chaitanya K. Jaladanki, Achal Ajeet Rayakar, Yap Xiu Huan, Hao Fan

## Abstract

Acetylcholinesterase (AChE) inhibition is a key mechanism in the treatment of neurodegenerative diseases and in counteracting toxic exposures to pesticides and nerve agents. However, virtual screening of AChE remains challenging due to the enzyme’s structural flexibility and the chemical diversity of its covalently binding inhibitors. In this study, we developed an in silico protocol that integrates multi-structure covalent docking and machine learning (ML) consensus scoring to improve the prediction of AChE inhibitors. We analyzed 65 ligand-bound (holo) human AChE crystal structures using hierarchical clustering to identify four representative conformations, along with one high-resolution apo structure, for multi-structure docking. A curated library of 412 organophosphate and carbamate inhibitors was then docked covalently and non-covalently into each receptor conformation. The resulting docking scores were evaluated against inhibitors’ experimental logIC_50_ values using Spearman’s rank correlation coefficient (r). Covalent docking outperformed non-covalent docking (r values up to 0.54 vs 0.18), and our ML consensus model trained on the five structures’ covalent docking scores achieved the highest predictive accuracy (r = 0.70), surpassing all single-structure and conventional consensus baselines. Chemical cluster analysis revealed structure–activity trends based on ligand flexibility, polarity, and aromaticity. SHapley Additive exPlanations analysis highlighted the ML consensus model’s ability to flexibly distribute the influence each structure’s scores played on its predictions. It identified and exploited relationships based on its training dataset that would be difficult to anticipate through a manual analysis of individual structures’ docking performance metrics. This framework is broadly applicable to other covalently targeted proteins, offering a generalizable and interpretable strategy for data-driven covalent inhibitor discovery.

## Introduction

Acetylcholinesterase (AChE) is a key enzyme responsible for hydrolyzing the neurotransmitter acetylcholine at cholinergic synapses in the central and peripheral nervous systems. The enzyme’s catalytic efficiency and essential role in neurotransmission make it a critical target for both therapeutic agents – used in the management of Alzheimer’s disease, Parkinson’s disease, myasthenia gravis, and other neurodegenerative disorders – and toxic compounds, such as organophosphates and carbamates, which act as irreversible inhibitors [1–6]. AChE inhibition leads to an accumulation of acetylcholine, resulting in overstimulation of cholinergic receptors and a cholinergic crisis. Owing to its well-characterized structure and pharmacological relevance, predicting the inhibition of AChE is critical for developing effective antidotes and therapeutic interventions [6–9].

AChE inhibition involves a complex two-step covalent modification process, as depicted in Figure 1. Initially, the substrate binds to the active site to form an enzyme–substrate complex. This is followed by an intermediate step in which the labile group is released, yielding a stable covalent enzyme–inhibitor complex. The energy changes associated with this reaction dictate the potency of the chemical, influencing its toxicity. Given the complexity of AChE inhibition and the challenges of experimental testing, computational methods can help unravel its intricacies and enhance our understanding of toxicity mechanisms. Molecular modeling has proven to be a powerful tool for investigating protein-ligand interactions at the atomic level. Traditional Quantitative Structure-Activity Relationship (QSAR) approaches have been used to predict AChE inhibition [10–23]; however, these methods face limitations when applied to chemicals that deviate from the structural scaffolds of known AChE inhibitors. Structure-based molecular docking and virtual screening, which use three-dimensional structures of target proteins, provide more accurate insights into molecular interactions. Nevertheless, most benchmarking studies of docking protocols like AutoDock Vina [24], GOLD [25], and Glide [26] have been performed using single crystallographic structures of AChE to evaluate their ability to reproduce known binding conformations and enrich active compounds in virtual screening [27–33]. These studies reveal that certain protocols, notably GOLD with the ChemPLP scoring function [30], demonstrate high reliability and enrichment factors when benchmarked against a single protein structure. However, their focus on non-covalent interactions and static conformations may limit their applicability for covalently binding inhibitors such as organophosphates and carbamates especially given the structural diversity of both the AChE active site and its covalent inhibitors.

**Figure 1.**
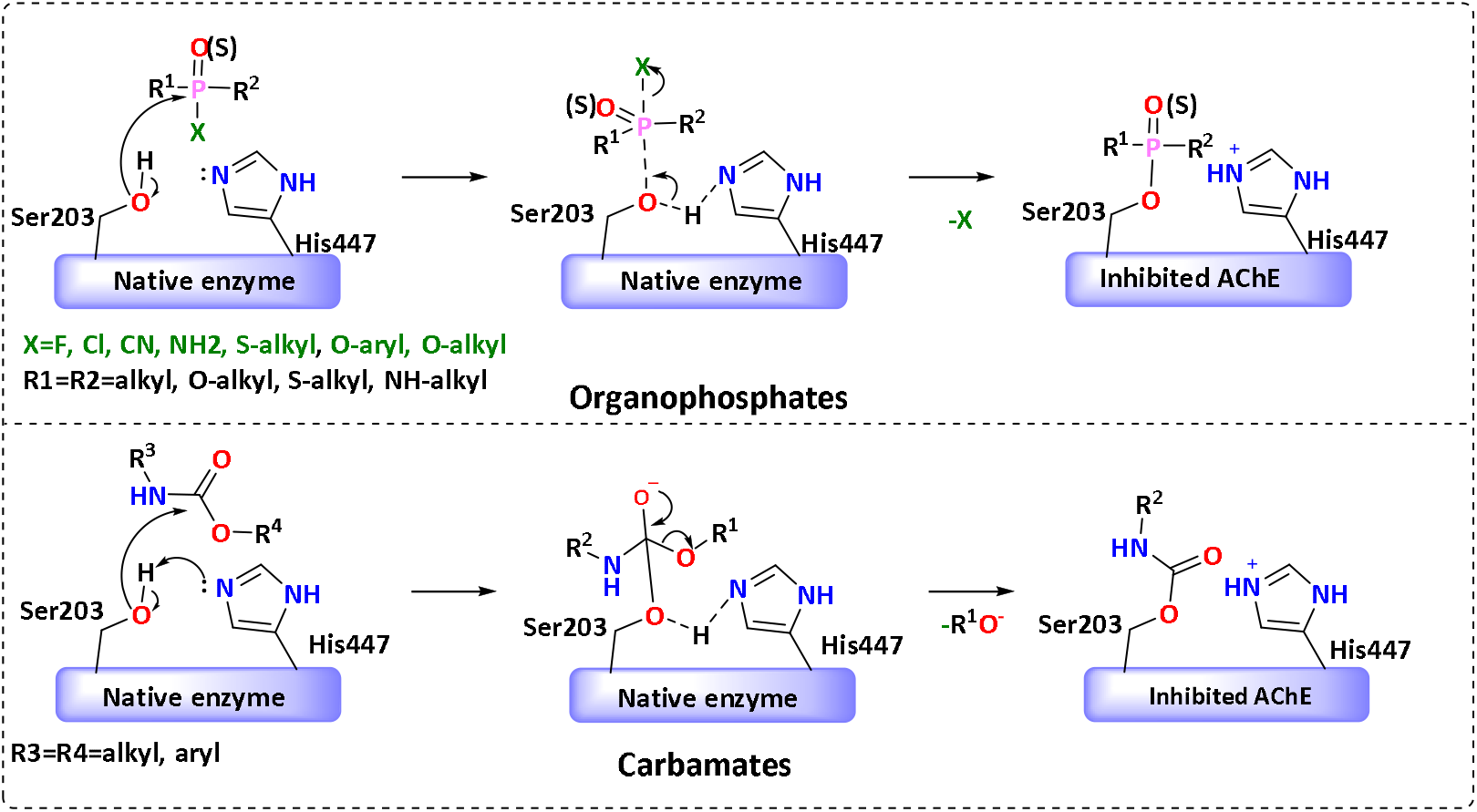
Biochemical reactions between organophosphates and carbamates and acetylcholinesterase (AChE).

To address the potential limitations of single-structure, non-covalent docking protocols, this study introduces an integrated in silico framework that incorporates protein structure clustering, multi-structure covalent docking, and machine learning (ML) consensus scoring (Figure 2) to enhance the prioritization of covalent inhibitors by predicted potency against AChE. While ML consensus scoring has been successfully applied in non⍰covalent virtual screening, most notably to consolidate docking protocol outputs across multiple scoring functions, protein conformations, or binding poses [34–37]. However, such approaches have typically excluded covalent docking protocols – even in studies ultimately focused on covalent inhibitors (e.g., TMPRSS2) [38] – with recent ML consensus pipelines often relying on non⍰covalent docking outputs (e.g., β⍰lactamase) [39]. In particular, multi⍰structure docking strategies have demonstrated improved predictive performance in non⍰covalent settings [34,35], but their application to covalently binding ligands remains limited [38–40]. To our knowledge, this study presents the first ML consensus model trained directly with covalent docking scores from multiple structures, explicitly leveraging both pre⍰and post⍰reaction complex energies, to yield a unified potency estimate for each compound.

**Figure 2.**
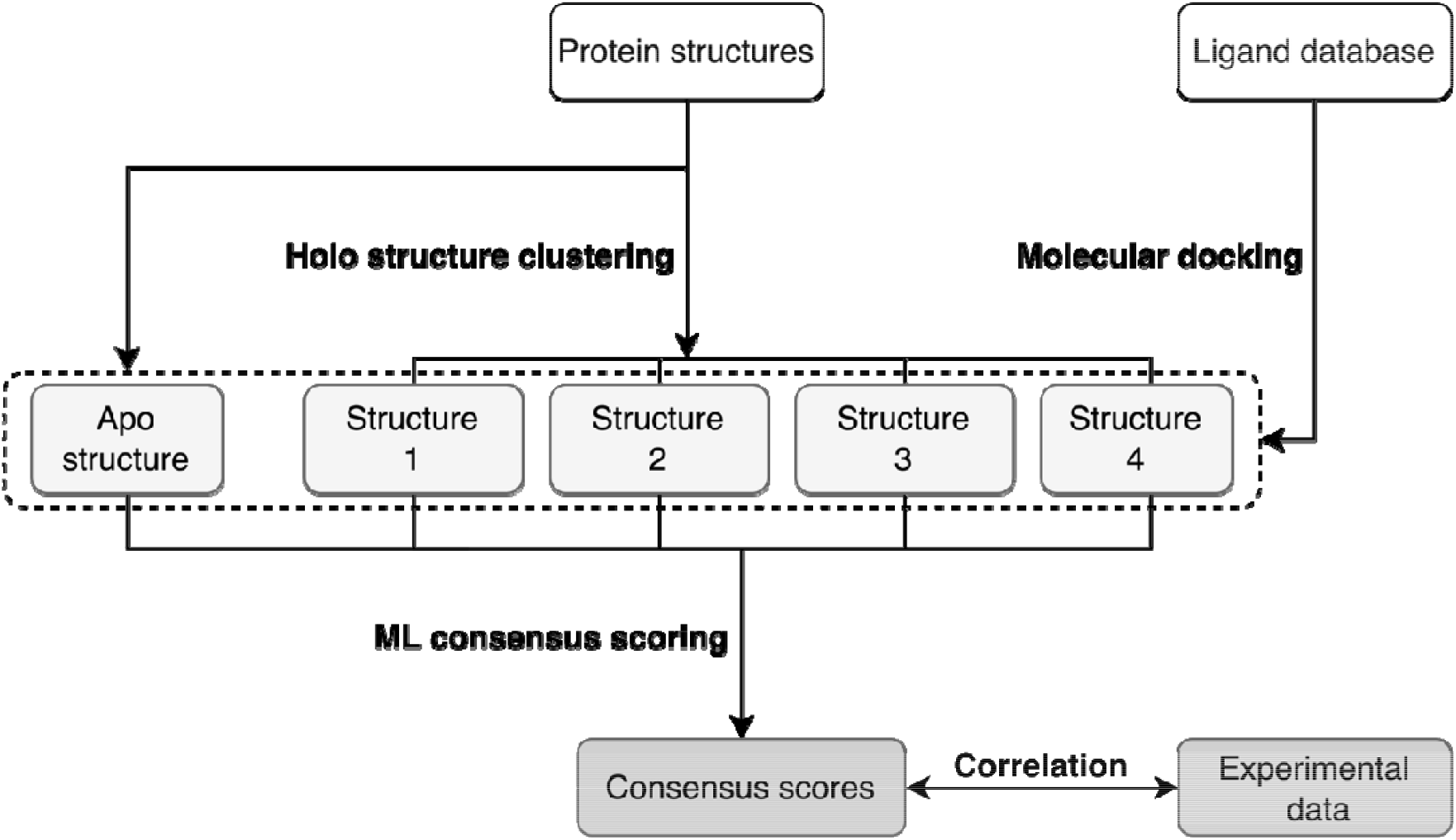
Workflow for multi-structure docking-based virtual screening protocol for predicting binding affinities of AChE inhibitors.

Empirically, this model surpasses both single-structure and heuristic consensus scoring approaches in terms of rank correlation, more accurately prioritizing the most potent inhibitors. In parallel, we performed a structure– activity analysis across chemically diverse clusters of covalent AChE inhibitors to contextualize model performance and clarify how receptor conformation and ligand properties jointly influence docking accuracy. Although demonstrated here for AChE, our strategy, which integrates multi-structure covalent docking with data-driven consensus modeling, offers a broadly applicable template for virtual screening of other targets where covalent binding and receptor flexibility challenge conventional approaches.

## Results and discussion

Our aim was to develop and evaluate a computational protocol for predicting human AChE inhibitors. To incorporate protein flexibility, we clustered 65 human AChE holo structures (80 chains) and selected four representative conformations for molecular docking, along with an additional apo structure. A curated library of 412 organophosphate and carbamate inhibitors was screened using both non-covalent and covalent docking approaches against these five representative structures. Predictive performance was then assessed by correlating docking scores with experimental logIC_50_ values in terms of Spearman’s rank correlation (r). Spearman’s rank correlation is a non-parametric measure that evaluates how well the relationship between two variables can be described by a monotonic function, making it well suited for tasks focused on prioritization rather than precise numerical prediction. This rank-based optimization criterion reflects the practical objective of docking: to prioritize compounds by their relative binding affinities rather than to predict absolute potency with high precision.

### Clustering Analysis of AChE Structures

To capture the structural diversity of AChE, hierarchical and k-means clustering were performed using the root mean square deviation (RMSD) of 25 binding-site residues (Figure S1). This clustering analysis of 80 individual protein chains from 65 holo structures (as detailed in Methods and Materials) identified four distinct structural clusters (Table 1). Cluster 1 was the largest, comprising 56 and 49 structures in hierarchical and k-means clustering, respectively, with 48 structures common to both methods. Cluster 2 comprised 12 structures in hierarchical clustering and 20 in k-means, with 11 structures common to both. Clusters 3 and 4 each contained six structures in hierarchical clustering, while k-means identified six and five structures, respectively; in both cases, five structures overlapped. The centroid structure from each cluster, as identified by hierarchical clustering, was selected for docking. These included AChE structures bound to a non-covalent ligand (PDB ID: 4M0E), a covalent organophosphate inhibitor (PDB ID: 6NTO), a covalent organophosphate inhibitor with a co-crystallized oxime molecule (PDB ID: 6WUY), and a metallosupramolecular inhibitor (PDB ID: 8AEN). In addition, the apo structure with the best resolution (PDB ID: 1B41) was included. These representative structures allowed a comprehensive docking analysis that accounted for protein flexibility and ligand-binding variations.

**Table 1.** Summary of clustering analysis performed on 80 AChE protein chains extracted from 65 holo crystal structures.

| Cluster | Hierarchical clustering | K-means clustering | Common structures |
| --- | --- | --- | --- |
| 1 | 56 | 49 | 48 |
| 2 | 12 | 20 | 11 |
| 3 | 6 | 6 | 5 |
| 4 | 6 | 5 | 5 |

The crystal structures of human AChE feature a deep, narrow catalytic gorge comprising two key regions: the catalytic active site (CAS), located at the bottom of the gorge, and the peripheral anionic site (PAS), situated near the gorge entrance. The CAS includes the catalytic triad (S203, H447, E334), catalytic anionic site residues (W86, Y133, F338), the acyl pocket (F295, R296, F297), and the oxyanion hole, which is formed by the backbone amides of G121, G122, and A204 and plays a critical role in stabilizing the transition state during catalysis. The PAS consists of residues Y72, D74, T75, Y124, V294, W286, Y337, and Y341, which modulate ligand access and orientation within the gorge. To assess conformational variability across binding pockets, the five representative structures were structurally aligned using 4M0E as the reference. All five structures showed high conservation of the core binding pocket architecture, with backbone RMSD values below 1.2 Å. However, notable conformational differences were observed in both CAS and PAS regions upon ligand binding. Superimposing 6NTO onto 4M0E revealed subtle conformational changes in CAS residues E334 and H447, accommodating the covalent adduct formed between the organophosphate and S203. The oxyanion hole was partially occupied by the phosphoryl oxygen (P=O) of the covalently bound organophosphate, resulting in conformational changes in neighboring residues, including W286, F295, and R296 (Figure 3). Similarly, superimposing 6WUY onto 4M0E showed that CAS residues underwent changes similar to those in 6NTO, along with notable conformational changes in PAS residues including W286, Y337, and Y341. In 6WUY, the oxime-based reactivator HI-6 interacted with the PAS through a salt bridge with D74 and π– π stacking interactions with Y124, W286, and the reoriented Y337 and Y341 (Figure 3). The tight binding of HI-6 at the PAS induced conformational changes in PAS and subtle allosteric effects on CAS residues. By contrast, superimposing 8AEN onto 4M0E revealed distinct conformational changes in CAS and PAS regions, involving residues Y72, Y124, F295, F297, and W286 (Figure 3). The ligand in 8AEN occupied both the CAS and PAS, forming a network of π–π stacking and cation–π interactions. In the CAS, the ligand’s positively charged aminium group formed a cation–π interaction with W86, stabilizing the ligand within the active site. In the PAS, residues F295, F297, and Y341 reoriented to optimize π–π stacking with the ligand’s pyridine moiety. These changes collectively resulted in a narrowed active-site gorge, with tighter packing around the ligand.

**Figure 3.**
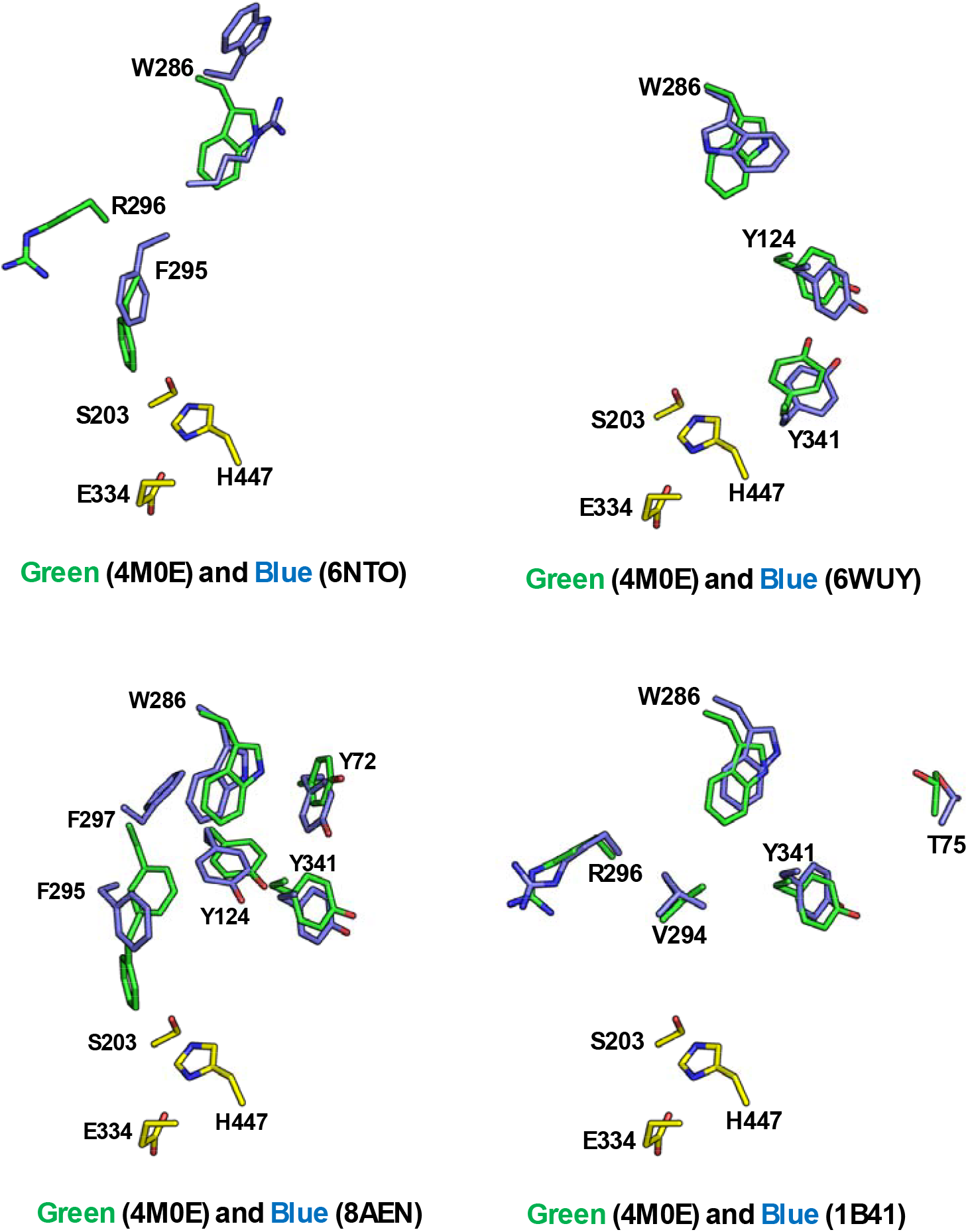
The cluster representative structures and apo structure of AChE are superimposed over 4M0E, highlighting key structural variations. (A) Superimposition of 6NTO over 4M0E, major structural changes observed in W286, F295, and R296 (B) Superimposition of 6WUY over 4M0E Major structural changes observed in Y124, W286, and Y341 (C) Superimposition of 8AEN over 4M0E, Major structural changes observed in Y72, Y124, F295, F297 and Y341 (D) Superimposition of 1B41 over 4M0E Major structural changes observed T75, W286, V294, R296, and Y341. The catalytic residues are highlighted in yellow (4M0E).

In addition, comparison of the apo structure 1B41 with 4M0E revealed that, in the absence of ligand, PAS residues such as T75, W286, V294, R296, Y337, and Y341 adopted more open conformations (Figure 3). Altogether, these observations demonstrate the structural flexibility of human AChE, which facilitates the binding of diverse inhibitors and underscores the importance of accounting for conformational diversity in docking studies to improve predictive accuracy.

### Analysis of Different Chemical Classes

To assess the chemical diversity among the covalent inhibitors, all compounds were clustered based on molecular fingerprint similarity, using a Tanimoto coefficient (T_c_) cutoff of 0.6. This approach yielded 12 chemical clusters, each representing a distinct molecular scaffold or pharmacophore class. The clustering approach aimed to capture structurally and functionally diverse chemotypes, potentially including phosphonates, carbamates and related analogues. Representative ligands from each cluster are illustrated in Figure 4 and Table S1. These representatives reflect the core features of their respective clusters. They were subsequently used for structure-based docking and incorporated into machine-learning models to evaluate structure–activity relationships and predictive performance across AChE conformations.

**Figure 4.**
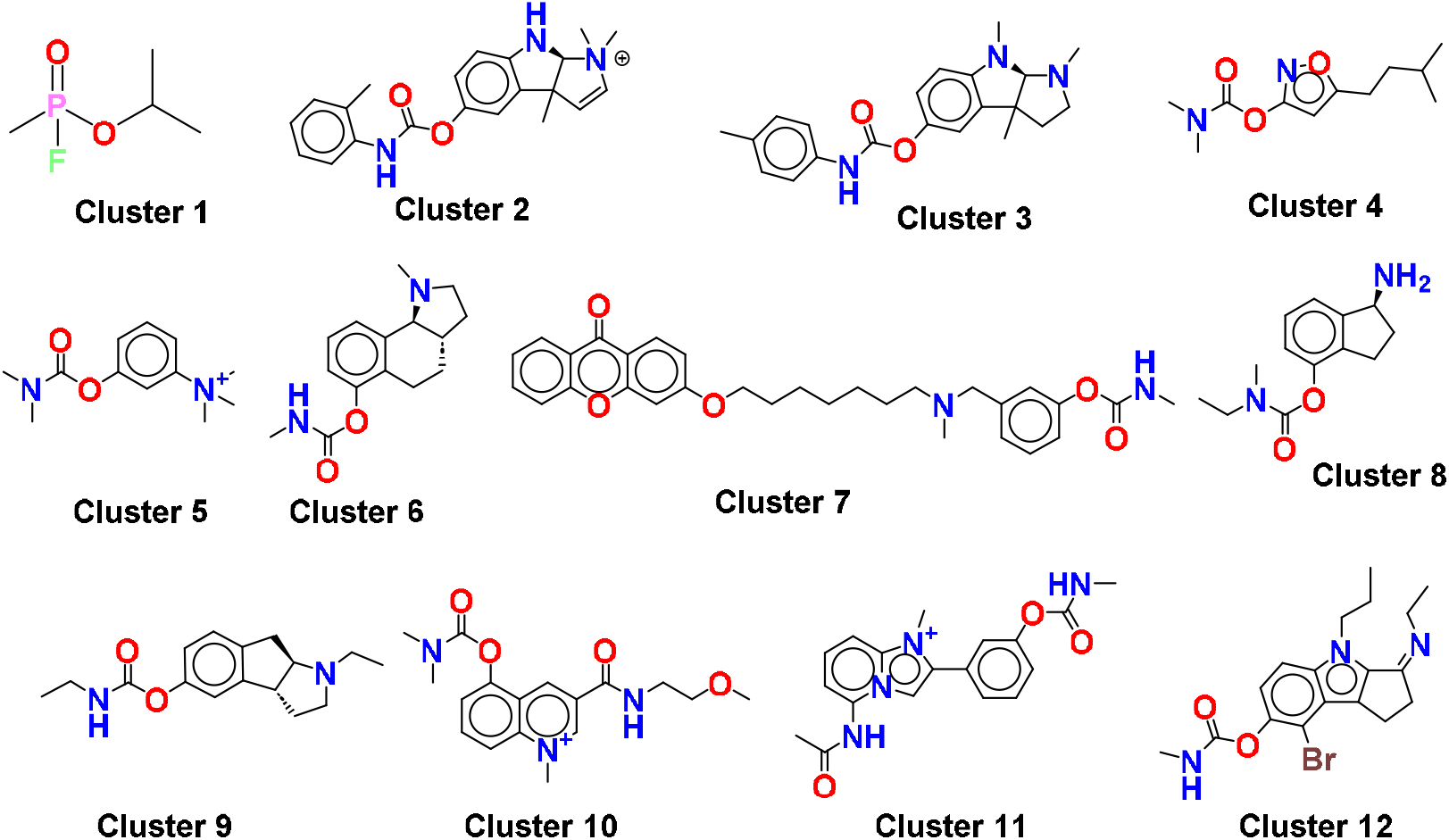
2D structures of representative chemicals from each of 12 clusters.

### Cognate ligand docking

To assess the reliability of our docking methods, we redocked the cognate ligands into their respective holo crystal structures. Non-covalent docking was employed for the non-covalent complex 4M0E, while covalent docking was used for the covalent complexes (6NTO, 6WUY, and 8AEN). Prior to docking, all bound ligands were removed from the crystal structures. Docking accuracy was evaluated by calculating the RMSD between predicted poses and the corresponding crystallographic ligand structures. The RMSD values were 0.9 Å for 4M0E, 1.8 Å for 6NTO, 1.6 Å for 6WUY, and 1.8 Å for 8AEN – all below the 2 Å threshold – which indicates reliable pose reproduction by both non-covalent and covalent docking methods (Supplementary Figure S3). The relatively low RMSD for 4M0E can be attributed to the ligand dihydrotanshinone I, which has a rigid, aromatic structure with only a single rotatable bond. In contrast, the slightly higher RMSD values for 6NTO, 6WUY, and 8AEN likely reflect the complexities introduced by covalent bond formation between the ligands and the enzyme, as well as the subsequent energy minimization of the ligands and active-site residues during covalent docking. These results confirm that our docking protocols reliably reproduce experimentally observed binding modes, supporting their use in subsequent virtual screening studies.

### Virtual screening and ML consensus scoring

We virtually screened the library of 412 covalent ligands, organized into 12 clusters, against five representative conformations of AChE. Both non-covalent and covalent docking protocols were applied: the former employed flexible-ligand docking, while the latter incorporated bond-formation constraints to model irreversible inhibitor binding. With these docking scores, we compared a range of single- and multi-structure docking strategies. Specifically, both kinds of strategies were considered in terms of four (one non-covalent and three covalent) docking score configurations. The covalent configurations encompassed using the pre- and post-reaction complex docking scores both individually and together. Under each configuration, docking strategies were compared in terms of the Spearman’s rank correlation (r) achieved between their outputs and experimental logIC_50_ values.

Because multi-structure docking produces multiple scores for the same compound, a consensus method was required to consolidate them. We therefore evaluated three approaches: (i) the mean consensus, which averages all input scores; (ii) the minimum consensus, which selects the most favorable (lowest) score; and (iii) the ML consensus, which applies a supervised machine learning model to integrate docking scores while capturing latent relationships among them.

Specifically, the ML consensus introduced within this work employs Random Forest regression^33^, which uses an ensemble of decision trees to create its predictions. Under each input docking score configuration, the ML consensus’ Random Forest was trained on a fixed set of 226 compounds randomly sampled from the 412 under consideration. The remaining 186 compounds, representing approximately 45% of the total, were held out for testing in comparison with other methods. This sampling was stratified by compound cluster to ensure the 12 clusters were equivalently represented across the training and testing compound sets. Moreover, to address the non-uniform distribution of compound potencies across the data (Figure S2), the training procedure incorporated a weighting scheme based on kernel density estimation (KDE) of the logIC_50_ values. Compounds occupying sparsely populated regions of the activity spectrum (particularly the strongest binders) were assigned greater weight during training. This correction was intended to encourage the Random Forest model to more equitably learn structure– activity relationships across the full dynamic range of biological response.

Table 2 presents the Spearman’s rank correlation (r) values obtained for the single- and multi-structure docking methods under the non-covalent and three covalent docking score configurations. Broadly, methods performed comparably well or better with covalent docking scores than with non-covalent docking scores. Specifically, single-structure methods experienced a mean r gain of 0.18 points when using pre- and post-reaction complex docking scores together (in the form of the CovDock score, which is the mean of both scores) instead of non-covalent scores. The largest improvements were observed for structures 6WUY (Δr = 0.39) and 8AEN (Δr = 0.29). An exception to this trend was 1B41, which achieved consistently poor performance across docking score configurations, with r values of 0.13–0.16. This may be due to its unliganded state, which can result in a less optimized binding site, particularly for bulkier or more flexible compounds.

**Table 2.**
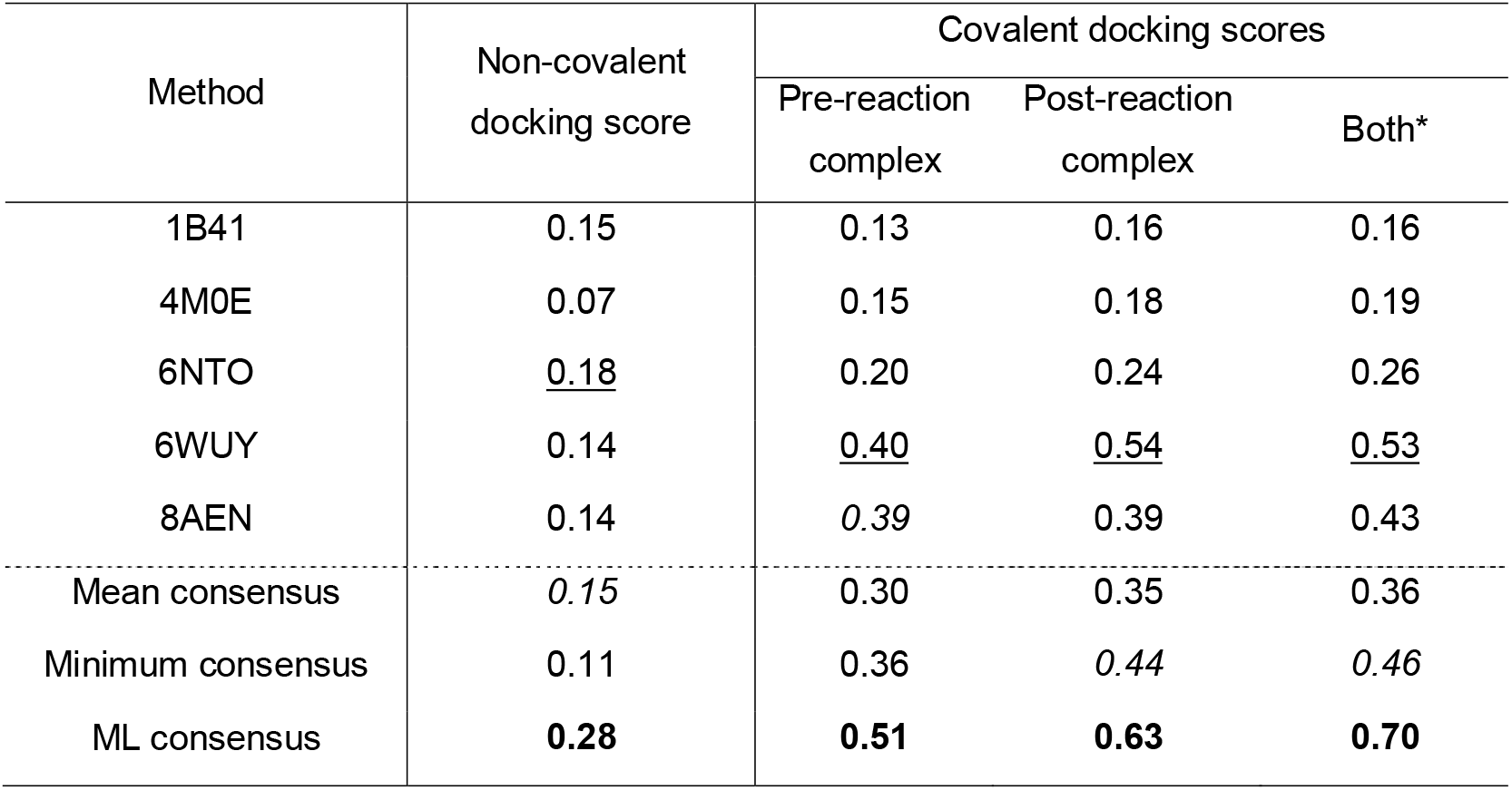
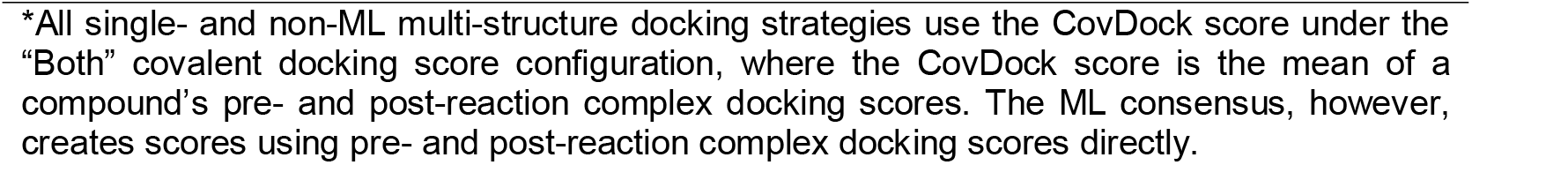
Spearman’s rank correlation coefficients between non-covalent and covalent docking scores (either single-structure or via consolidation across structures) and logIC_50_, taken over the 186 compounds not used in the ML consensus’ training. (**Best**, <u>second best</u>, *third best*).

| Method | Non-covalent docking score | Covalent docking scores |  |  |
| --- | --- | --- | --- | --- |
|  |  | Pre-reaction complex | Post-reaction complex | Both* |
| 1B41 | 0.15 | 0.13 | 0.16 | 0.16 |
| 4M0E | 0.07 | 0.15 | 0.18 | 0.19 |
| 6NTO | <u>0.18</u> | 0.20 | 0.24 | 0.26 |
| 6WUY | 0.14 | <u>0.40</u> | <u>0.54</u> | <u>0.53</u> |
| 8AEN | 0.14 | *0.39* | 0.39 | 0.43 |
| Mean consensus | *0.15* | 0.30 | 0.35 | 0.36 |
| Minimum consensus | 0.11 | 0.36 | *0.44* | *0.46* |
| ML consensus | **0.28** | **0.51** | **0.63** | **0.70** |

\*All single- and non-ML multi-structure docking strategies use the CovDock score under the “Both” covalent docking score configuration, where the CovDock score is the mean of a compound’s pre- and post-reaction complex docking scores. The ML consensus, however, creates scores using pre- and post-reaction complex docking scores directly.

**Table 3.**
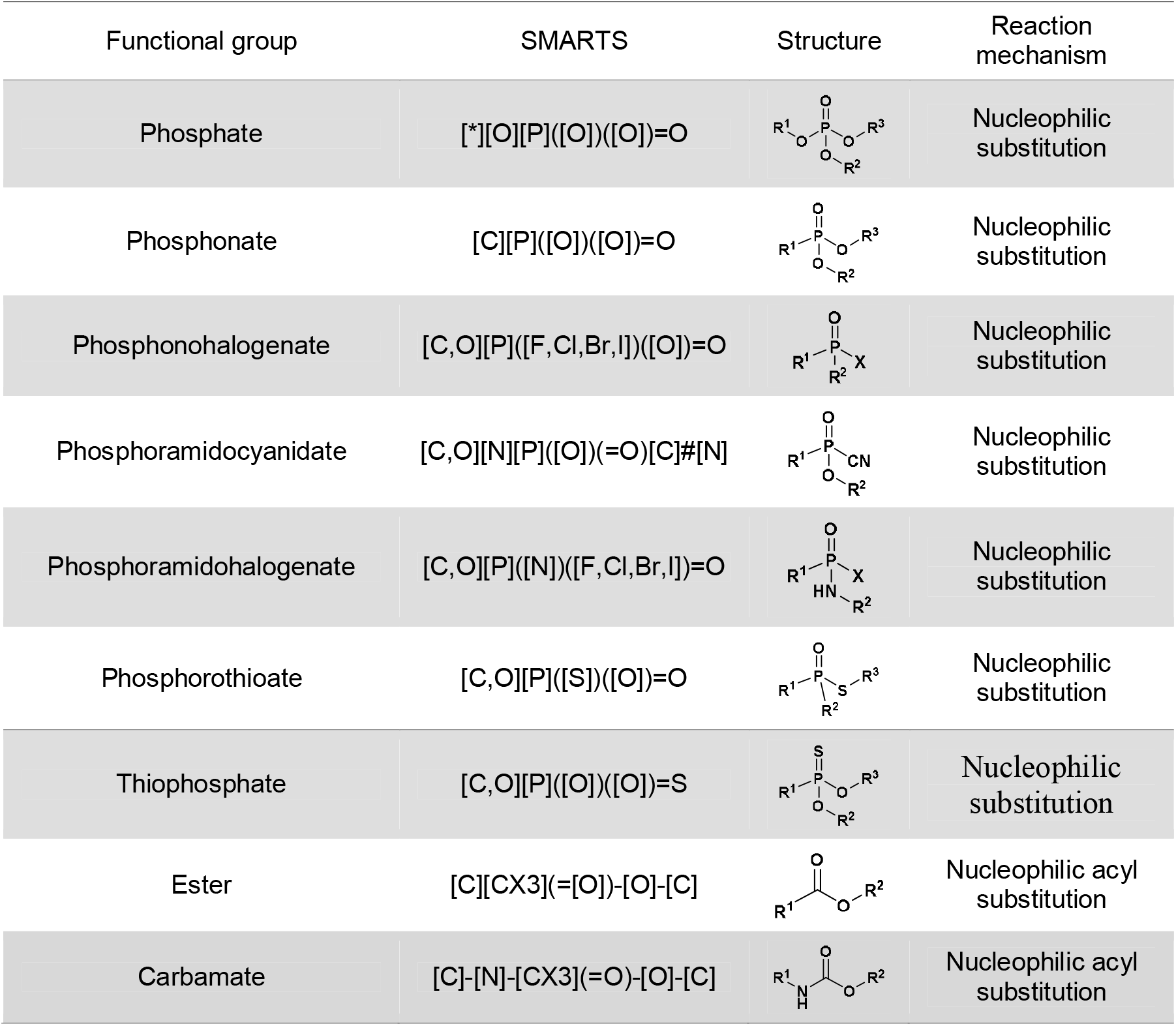
Covalent warheads used against AChE.

| Functional group | SMARTS | Structure | Reaction mechanism |
| --- | --- | --- | --- |
| Phosphate | [*][O][P]([O])([O])=O |  | Nucleophilic substitution |
| Phosphonate | [C][P]([O])([O])=O |  | Nucleophilic substitution |
| Phosphonohalogenate | [C,O][P]([F,Cl,Br,I])([O])=O |  | Nucleophilic substitution |
| Phosphoramidocyanidate | [C,O][N][P]([O])(=O)[C]#[N] |  | Nucleophilic substitution |
| Phosphoramidohalogenate | [C,O][P]([N])([F,Cl,Br,I])=O |  | Nucleophilic substitution |
| Phosphorothioate | [C,O][P]([S])([O])=O |  | Nucleophilic substitution |
| Thiophosphate | [C,O][P]([O])([O])=S |  | Nucleophilic substitution |
| Ester | [C][CX3](=[O])-[O]-[C] |  | Nucleophilic acyl substitution |
| Carbamate | [C]-[N]-[CX3](=O)-[O]-[C] |  | Nucleophilic acyl substitution |

Gains from using CovDock over non-covalent scores were also pronounced among the mean and minimum consensus methods, which improved by 0.21 and 0.35 r points, respectively. Notably, the mean and minimum consensus methods used compounds’ CovDock scores because they could not operate over the ten individual pre- and post-reaction complex docking scores, which are not directly comparable. The ML consensus, however, could use scores individually and achieved the highest overall performance, with an r of 0.70 – a 0.42-point gain over its performance with non-covalent docking scores. These substantial improvements demonstrated not only the advantage of covalent docking for ranking AChE inhibitor potency, but also the added value of processing covalent docking scores with the ML consensus method.

Most broadly, the ML consensus shows improvements of 0.10-0.17 r points over the best alternative consensus and single-structure methods across all docking score configurations. Importantly, the superior performance of the ML consensus when using both pre- and post-reaction complex docking scores compared to other methods that, by their design, can only use CovDock scores, is not due to the nature of the CovDock score itself. A revised, dynamic form of the CovDock score, which weighted the average of pre- and post-reaction complex docking scores according to the training dataset, did not achieve a higher r. This was true despite the weighting typically favoring the more informative post-reaction score (see Materials and Methods and Table S2). Notably, post-reaction docking scores exhibited greater predictive power than pre-reaction scores alone.

These findings highlight the critical contribution of post-reaction complex geometry in capturing key steric and electronic interactions arising from covalent bond formation, which directly govern inhibitor potency. By incorporating the structural consequences of bond formation, post-reaction docking scores more accurately reflected the biologically relevant final binding state that drives activity. This enhanced structural fidelity likely underpinned the improved predictive performance and aligned with biochemical principles, where covalent bond-induced conformational changes critically influence binding affinity.

Figure 5 illustrates the performance of the ML consensus within covalent configurations in greater detail. As shown in Figure 5(A), moving from pre-reaction to post-reaction complex docking scores, and then to using both together, reduced scatter around the line of best fit, complementing the increase in r. Figure 5(B) also complements correlation-based evaluations by illustrating compounds’ rank errors. Across every covalent docking score configuration, the ML consensus exhibited the narrowest, most sharply-peaked error distribution, with standard deviations that are 7.62-14.29 ranks lower than those of the minimum consensus and 5.33-10.35 ranks lower than those of the best single-structure method, docking to 6WUY. The tighter spread implied not just greater accuracy but, particularly fewer catastrophic mis-rankings – instances in which highly active inhibitors are relegated far down the ranked testing dataset or weak inhibitors and non-binders are promoted to the top. Figure 5(C) compares the ML consensus to the best-performing alternative method within each compound cluster, evaluating performance by cluster-r. Across all covalent docking score configurations (and particularly for those using post-reaction complex docking scores), the ML consensus achieved the largest number of best performances, outdoing its counterparts at the subpopulation level. Cluster-r captures how well methods preserve the relative potency ordering of compounds within individual clusters, in contrast to (overall) r which evaluates ranking performance across the entire testing dataset. Despite being trained to predict every compound’s potency, the ML consensus’ superior cluster-level ranking suggested that it was not only effective globally but also adept at preserving local structure–activity relationships.

**Figure 5.**
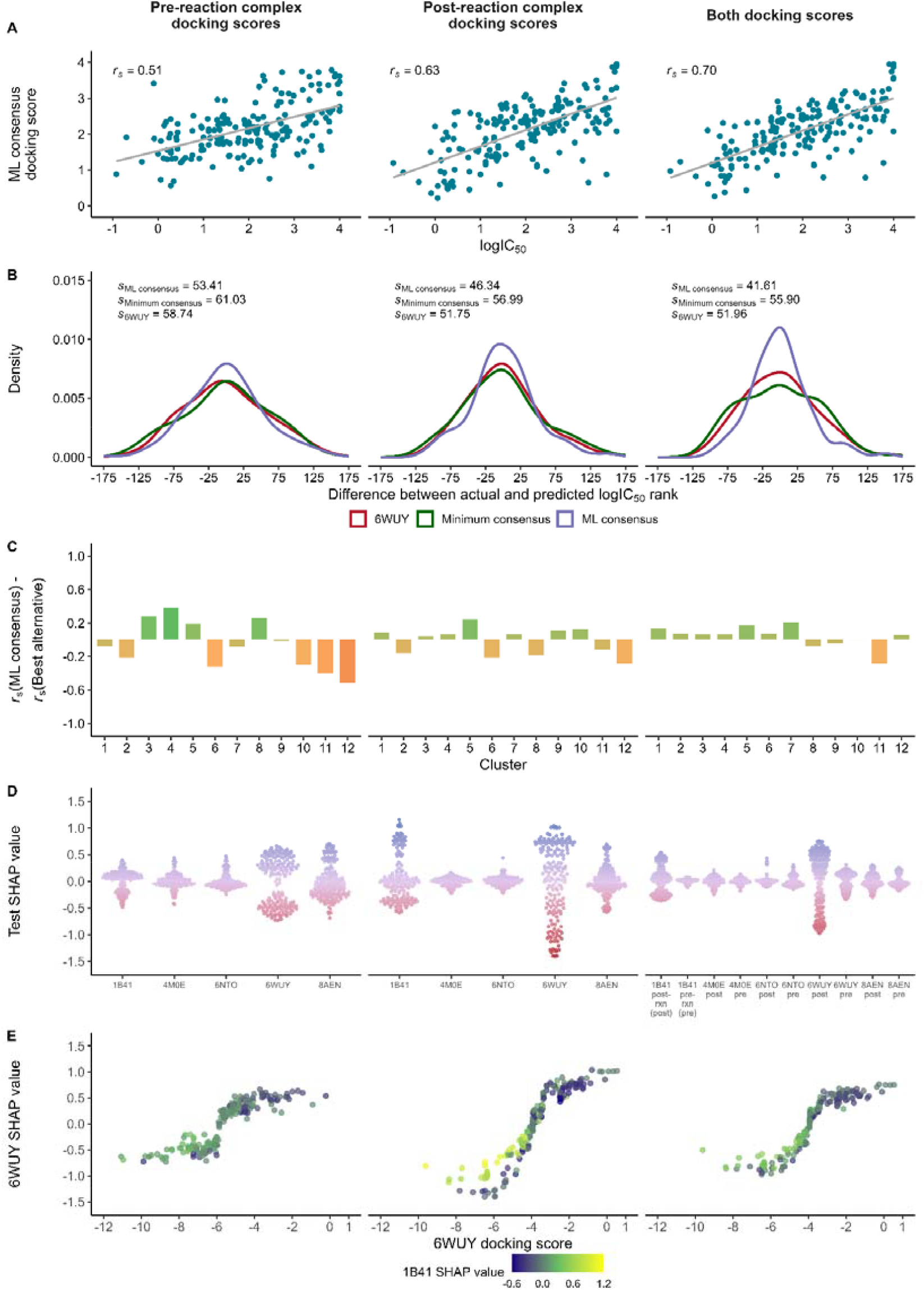
Enumeration and diagnostic of the ML consensus’s performance across each covalent docking score configuration. (A) Scatterplots of the consensus docking score versus logIC_50_. The ML consensus achieves better performance using both pre- and post-reaction complex docking scores than it does using either type alone. (B) Distributions of ranking errors for the ML-consensus, minimum-consensus, and 6WUY-alone methods, showing the ML consensus to have the smallest rank error spread. (C) The difference in the compound cluster-r achieved by the ML consensus and that achieved by the best alternative method. The best alternative varies by cluster. (D) Beeswarm plots of the testing SHAP values associated by each input docking score to the ML consensus. Each point in the beeswarm plot represents the docking score’s contribution for an individual testing compound. (E) Scatterplot of 6WUY docking scores’ SHAP values against the docking scores themselves, colored by the corresponding 1B41 SHAP value. Across all docking score configurations, the former relationship is approximately sigmoidal.

Figure 5(D) presents a mechanistic diagnostic of the ML consensus through SHAP (SHapley Additive exPlanations) analysis.^34,35^ Across all covalent docking score configurations, the ML consensus used scores in a highly selective manner: a small subset of structures contributed prominently, with broad, high-magnitude SHAP value distributions, while the remaining structures exhibited sharply narrower spreads centered near zero, indicating limited influence. Specifically, the docking scores of 6WUY and 8AEN contributed greatly to the ML consensus’ outputs, corresponding with the leading performances of the single-structure methods employing them. However, counterintuitively, 1B41’s docking scores played a prominent role within the post-reaction and CovDock configurations despite achieving the lowest standalone r values. This suggested that the ML consensus found 1B41 to contribute a conditionally useful signal.

Figure 5(E) therefore examines structures’ contributive relationships within the ML consensus in greater detail by plotting 6WUY’s SHAP values against its docking scores, with points colored by 1B41’s corresponding SHAP values. A sigmoidal pattern was observed across all covalent docking score configurations: 6WUY’s SHAP values increased with their docking scores but began to plateau at both extremes of the score range, suggesting that the model limited the marginal impact of very strong or very weak scores. Superimposed on this tempering relationship was a pattern in 1B41’s SHAP values: in regions where 6WUY’s SHAP values began to plateau – particularly for compounds with the most favorable (i.e., lowest) docking scores -- 1B41’s SHAP values were often of the opposite sign. While the association was not strictly inverse, it suggested that 1B41 also played a role in counterbalancing 6WUY’s contributions, but in a dynamic, context-sensitive manner beyond the observed sigmoidal relationship between 6WUY’s SHAP values and docking scores. Taken together, these patterns illustrated the ML consensus model’s ability to flexibly distribute influence across structures, identifying and exploiting relationships from its training dataset in a manner difficult to anticipate through conventional manual consensus rules and individual performance metrics alone.

Notably, the best non-ML consensus method, the minimum consensus, lagged behind the individual use of 6WUY across the three covalent docking score configurations by 0.04–0.10 points; however, it was either comparable to or considerably better than every other single-structure method. These results suggested that, in the common prospective discovery scenario where no target-specific activity data exist to gauge the reliability of individual structures, taking the minimum docking score across structures may serve as a conservative yet empirically robust hedge. This approach mitigates the risk of a poorly chosen structure and delivers a ranking performance that rivals the best single-structure alternative without requiring prior calibration. Nonetheless, we acknowledge that this empirical observation was only a preliminary indication, and that rigorous validation – via resampling, cross-cluster holdouts, and multi-target benchmarks – will be essential to establish minimum-consensus CovDock scoring as a generally reliable, calibration-free strategy. Moreover, because our primary objective was to maximize potency ranking accuracy for AChE – where ample historical activity data enabled supervised learning – the spotlight of this study remained on the ML consensus. Its capacity to exploit those data and systematically outstrip all heuristic aggregations underscored its practical value for targets with existing calibrating information.

### Chemical cluster correlation analysis

While SHAP-based analyses provide insight into how the ML consensus model dynamically used information from different receptor structures, they remained inherently limited to structure-wide docking signals. The model did not incorporate molecular interaction features or physicochemical descriptors as inputs, and we therefore could not directly attribute performance to specific ligand substructures or binding-site interactions. To further investigate the biophysical underpinnings of docking performance, we conducted an independent cluster-level diagnostic that examined how individual receptor conformations performed across the 12 chemically distinct compound groups. This finer-grained analysis offered complementary insight into the structural and chemical factors that shaped docking accuracy across the conformational landscape of AChE.

Specifically, compounds from these 12 chemical clusters exhibited disparate docking performances among the five AChE structures, influenced by properties such as molecular weight, flexibility, aromaticity, and charge (Figure 6). The clusters could be broadly separated into two mechanistic classes: organophosphates (Cluster 1) and carbamates (Clusters 2–12). Cluster 1 compounds consistently showed poor correlations between their docking scores and experimental logIC50, despite their high experimental potency. This discrepancy reflected the limitations of docking algorithms, which often failed to fully capture covalent binding contributions such as leaving-group lability and bond energies [41]. For example, organophosphates with fluoride (P–F) or cyanide (P–CN) leaving groups had low bond-dissociation energies (<80 kcal/mol; Table S3) compared with those of other leaving groups (e.g., S–CH3, O-phenyl) and with the ester bond of carbamates (>90 kcal/mol). This enabled their faster, irreversible covalent bond formation with the catalytic serine residue. CovDock evaluated only the final complex geometry and did not consider kinetic or thermodynamic factors that governed reactivity. Furthermore, the phosphorylated adducts formed during inhibition were resonance-stabilized and featured hypervalent phosphorus centers [42], characteristics inadequately captured by standard docking scoring functions. Moreover, the covalent docking protocol employed lacked reactivity-based descriptors (e.g., Fukui indices or electronic properties) that reflected the electrophilic potential crucial for modeling covalent inhibition.

**Figure 6.**
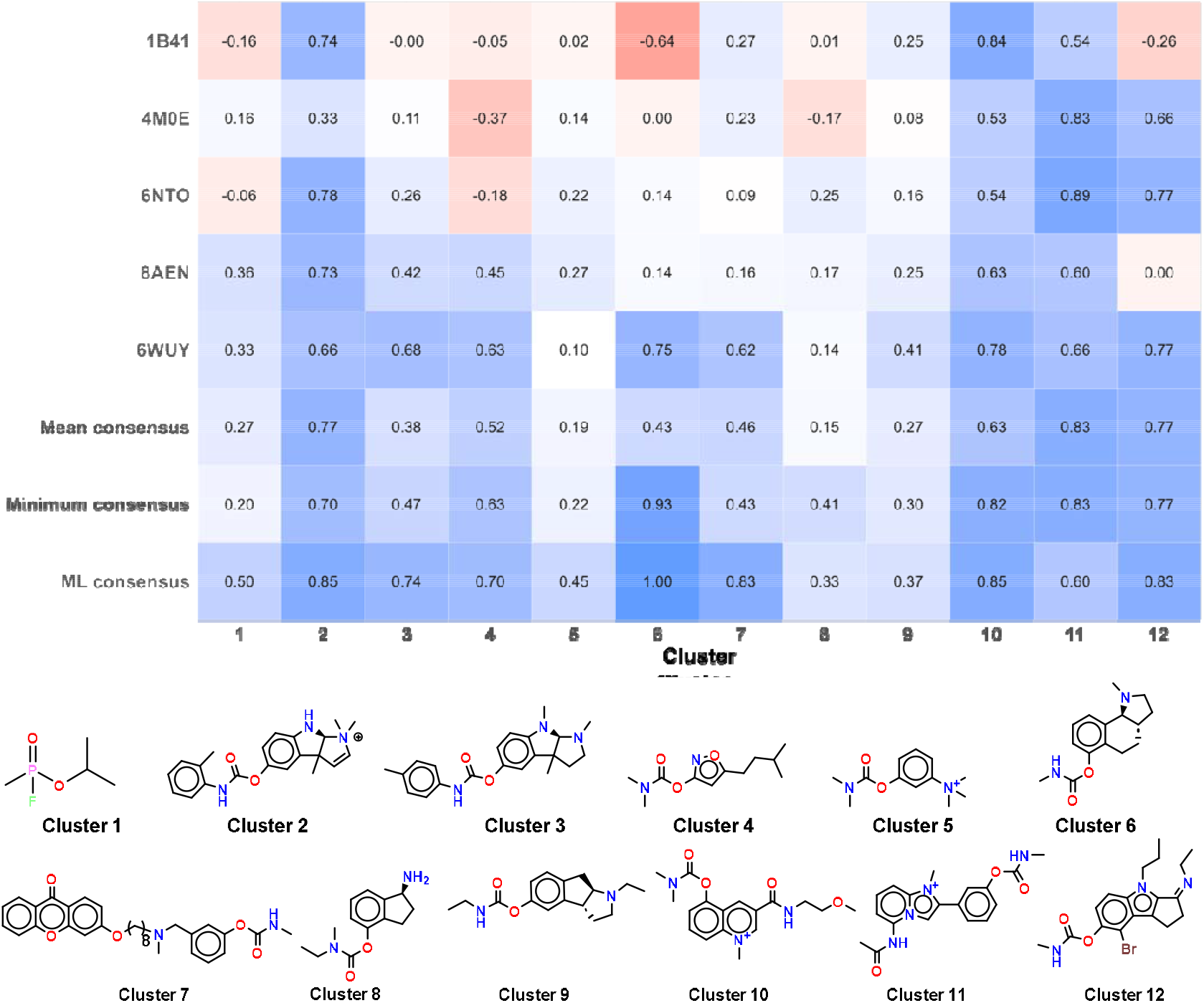
Heatmap of Spearman’s rank correlation coefficients (r) between predicted docking scores and experimental logIC_50_ values across all chemical clusters, with each cluster’s representative 2D compound structure. The columns represent individual chemical clusters, and the rows correspond to five AChE structures and three consensus scoring methods (mean, minimum, and ML consensus). Blue indicates a strong positive correlation, while red indicates weak or poor correlation.

The remaining 11 carbamate clusters inhibited AChE through reversible, non-covalent interactions. Their docking scores correlated more strongly with experimental activity than those of organophosphates, with an ML consensus r of 0.72 across clusters. Cluster 2 compounds, characterized by their planar aromatic rings (Figure S3A, B) and cationic groups, showed strong correlations between their docking scores and experimental potencies in four of the five AChE structures. This performance was facilitated by consistent π– π and cation–π interactions with key CAS residues W86 and Y337 (observed in 94% of the compounds in pre-reaction docking complexes) and by a favorable topological charge distribution (Figure S3C) that enhanced electrostatic complementarity with the binding site. These favorable charge-distribution and geometry features helped the docking program accurately predict their binding behavior. Similarly, Clusters 10–12, which contained biaryl and triaryl scaffolds, exhibited comparable interaction patterns and strong correlations.

Cluster 3 compounds (rigid, sp^3^-rich tricyclics) performed well only with open binding-site structures such as 6WUY (46% of compounds interacted with W86/Y337) but poorly otherwise (e.g., only 27% in 8AEN). These disparate performances may have been influenced by the compounds’ low aromaticity and numerous ring sp^3^ carbons (Figure S3D), which reduced stacking interactions and hindered optimal alignment, resulting in poorer docking accuracy. Similarly, Cluster 9 compounds (sp^3^-rich polycyclic scaffolds) also showed mediocre correlations in open binding-site structures, and performed even more poorly with narrower binding-site conformations. This reflected their limited conformational adaptability and reduced π-surface area.

Docking of Cluster 4, 5, and 8 compounds underperformed due to their low molecular weight (Figure S3E), moderate polarity, and limited hydrogen-bonding capacity, which reduced the number and strength of interactions they could form in the binding site. Clusters 4 and 5 also featured smaller aromatic groups (Figure S3A, B), leading to fewer and weaker π–π stacking interactions and limited hydrophobic and electrostatic complementarity, which often caused docking scores to underestimate relative potency. Docking of Cluster 6 compounds performed poorest overall (except in 6WUY). These compounds were bulky, less aromatic (low π-surface area, PISA; Figure S3G), and built around structurally rigid core scaffolds with few rotatable bonds, higher sp^3^ carbon counts, and a high degree of ring fusion (Figure S3A, D, E). These features restricted conformational adaptability, diminished π-π stacking, and hindered accommodation within narrower active-site gorges during docking. Their low polarity (polar surface area, TPSA ≈ 3 Å^2^) and absence of hydrogen-bond donors further weakened interactions in hydrophilic sub-pockets, reducing docking accuracy. Cluster 7 compounds, characterized by their long flexible alkyl linkers (>C10, Figure S3H), also exhibited poor correlations, except when docked in 6WUY. Their linker flexibility favored PAS contacts in the more open conformation but also prevented stable interactions deep in the catalytic gorge, which reduced docking accuracy.

In sum, docking performance varied substantially across both AChE structures and ligand clusters. The 6WUY structure (co-crystallized with an organophosphate and HI-6 oxime) performed best, with strong correlations in 8 of 12 clusters, likely reflecting ligand-induced flexibility in both CAS and PAS regions. Conversely, the apo structure 1B41 performed worst, showing reasonable correlation only for Clusters 2 and 10. From a clusterwise perspective, clusters composed of flat, aromatic, and charged compounds generally supported better docking, whereas rigid, bulky, or small non-aromatic compounds were less successful because the program struggled to fully account for shape, flexibility, and subtle non-covalent interactions. These results underscored the limitations of static docking for flexible or covalent ligands and emphasized the importance of selecting AChE conformations tailored to the chemical classes of interest in future virtual screening campaigns.

## Conclusion

In this study, we demonstrated that integrating multi-structure covalent docking with ML consensus scoring significantly enhanced the predictive accuracy of virtual screening for AChE inhibitors, with improvements in Spearman’s rank correlation of up to 0.27 points over the best-performing conventional alternatives. By leveraging five structurally distinct holo and apo AChE crystal structures, we showed that receptor flexibility played a key role in accurately ranking the binding affinities of chemically diverse covalent inhibitors, with individual structures exhibiting differential predictive power across inhibitor chemical clusters. Accordingly, the ML consensus model, trained on these structures’ covalent docking scores, consistently outperformed both single-structure and heuristic consensus approaches by selectively adjusting the contribution of each structure’s score within its consensus prediction.

Despite its strong performance for AChE, our strategy’s generalizability might be constrained for targets with limited structural data or different binding mechanisms. Nonetheless, and relatedly, our chemical cluster analysis revealed that ligand properties such as aromaticity, flexibility, charge, and electronic characteristics substantially impacted docking accuracy – factors not currently utilized by the ML consensus. Incorporating these and other physicochemical descriptors as auxiliary features could enhance its robustness for such targets.

Taken together, these findings established a compelling proof-of-concept for combining multi-structure covalent docking with ML consensus scoring in the context of a highly flexible, covalently targeted enzyme. This work offered a robust and extensible framework that could be readily adapted to other targets where induced fit, allosteric modulation, or covalent reactivity complicated traditional docking-based prioritization. Future applications to similarly challenging systems – such as proteases, kinases with reactive warheads, or epigenetic enzymes with flexible binding pockets – may further demonstrate the general utility of this strategy and accelerate the discovery of potent, selective covalent inhibitors in drug development.

## Key points

- We propose a proof-of-concept workflow that integrates protein structure clustering, multi-structure covalent docking and machine learning (ML) consensus scoring to rank covalent acetylcholinesterase (AChE) inhibitors.
- We construct an archetypal five-structure docking panel by clustering 65 AChE crystal structures and use it to generate pre- and post-reaction complex docking scores that a Random Forest consensus model learns from to rank inhibitor potency.
- On a held-out set of 186 organophosphate and carbamate inhibitors, the ML consensus attains a Spearman rank correlation of 0.70, exceeding the best single-structure and heuristic consensus alternatives by 0.16-0.24 points.
- This generalizable approach offers a template for applying multi-structure covalent docking with data-driven ML consensus scoring to other flexible proteins where single-structure docking may underperform.

## Methods and materials

### Protein database preparation

A total of 75 crystal structures were retrieved from the RCSB database [43], comprising both apo and holo forms. The dataset included 10 apo structures and 65 holo structures. The protein structures were prepared with Protein Preparation Wizard [44], including an optimization step of hydrogen bond assignments using ProtAssign from Schrödinger. Both chains of holo structures were used for clustering analysis to select representative protein structures for subsequent simulations. Additionally, one apo structure with the highest resolution was selected.

### Protein structure clustering

From an initial set of 65 AChE holo structures comprising 110 individual protein chains, 80 chains were selected for structural clustering based on two criteria: (i) inclusion of at least one representative chain per holo structure, and (ii) selection of chains exhibiting pairwise RMSD values greater than 1.0 Å within the binding site region. This filtering step was intended to maximize structural diversity while minimizing redundancy, thereby enhancing the representativeness of the clustering analysis. Each protein chain superposition was conducted based on the binding site residues side chains, defined as those within 5 Å of the ligand, totaling 25 cumulative residues (Figure S1). Hierarchical clustering [45] was performed using the atomic Root Mean Square Deviation (RMSD) of side chains of binding site residues. Hierarchical clustering generated four clusters with RMSD cutoff set at 1 Å (Table 1). Simultaneously, k-means clustering yielded four clusters (Table 1) [45].

A representative structure from each cluster was selected by choosing the structure with the highest number of neighboring structures, with an RMSD value below the specified threshold for subsequent docking analysis. The selected representative structures, along with a description of their bound ligands, were: 4M0E (non-covalent ligand), 6NTO (covalent phosphate ligand), 6WUY (covalent phosphate ligand with co-crystallized oxime), 8AEN (metallo supramolecule ligand).

### Compound database collection

A comprehensive ligand database was curated, comprising 19,875 compounds with activity data (IC_50_, EC_50_, K_i_, K_d_) from diverse sources such as ChEMBL, BindingDB, Drugbank [46], etc. After removing duplicates and filtration, 4,781 unique compounds with activity data below 10 µM were obtained. These compounds were further classified into covalent inhibitors, comprising 382 carbamate-based inhibitors and 30 organophosphate inhibitors. For subsequent docking studies, only covalent inhibitors were considered. The compounds are prepared by using LigPrep module of Schrodinger [47], The Epik module of Schrodinger was used to generate the ionization state at pH 7.0 ± 2.0, using the OPLS4 force field.

### Chemical compound clustering

Covalent AChE inhibitors were clustered using hierarchical clustering based on 64-bit Daylight-like molecular fingerprints. Pairwise Tanimoto similarity was calculated. Hierarchical clustering was performed using the average linkage method, with a Tanimoto similarity cutoff of 0.6 to define cluster boundaries. The resulting clusters were used to assess chemical diversity and ensure balanced representation in model training and testing.

### Covalent docking

Initially, we attempted to identify the covalent docking reaction indicating the catalytic serine – organophosphates or carbamate covalent interaction using Glide’s predefined (built-in) and custom covalent reactions available on Schrödinger’s covalent reactions repository. Despite conducting thorough search, this approach did not yield a suitable algorithm. Subsequently, we developed a customized Glide covalent docking [48] reaction algorithm for organophosphates or carbamates. The algorithm for organophosphates was designed to specify the electrophilic phosphorus center of organophosphates, the nucleophilic hydroxyl group of the serine residue (Ser-OH) on the target protein, ligand attachment atom on a receptor (O), receptor attachment atom on a ligand (P), formal charge (0), and bonds created (1 bond between an oxygen atom of Ser-OH and phosphorus atom of the phosphate group on organophosphate) and/or broken (1 bond on a ligand (P–X(halogen), P–OR, P–CN, P–S or P–N)) during a reaction process. The algorithm for carbamates was designed to specify the electrophilic carbonyl (C=O) center of carbamate, the nucleophilic hydroxyl group of the serine residue (Ser-OH) on the target protein, ligand attachment atom on a receptor (O), receptor attachment atom on a ligand (C=O), formal charge (0), and bonds created (1 bond between an oxygen atom of Ser-OH and carbon atom of the carbonyl on organophosphate) and/or broken (1 bond on a ligand [N]-(C=O)-[O]) during a reaction process. We conducted covalent docking using CovDock to assess the binding of 412 covalent inhibitors to four representative structures from each cluster and one apo structure of AChE. The docking simulations were performed in standard precision mode, utilizing the OPLS4 force field. The grid boxes were centered based on the bound ligand, and both inner and outer boxes had dimensions of 10 Å × 10 Å × 10 Å and 30 Å × 30 Å × 30 Å, respectively. The oxygen atom of the catalytic residue Ser203 was specified as the reaction site for the formation of a covalent bond, either with the carbamate or phosphate moiety of the selected inhibitor. Post-docking, a refinement of binding poses was executed with a minimization radius of 5.0 Å. The resulting docking poses were ranked based on both Prime energy and docking score. CovDock generates three types of scores for each compound: the pre-reaction complex score, the post-reaction complex score, and a composite CovDock score, defined as:

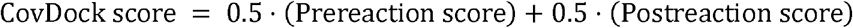

*Qikprop Analysis:* the QikProp module of Schrödinger was used to calculate physicochemical properties of the molecules. it predicts the Absorption, Distribution, Metabolism, Elimination (ADME) properties and bioavailability of the compounds.

All the quantum chemical calculations were conducted using the Jaguar module [49]. Geometry optimizations and frequency calculations were performed using density functional theory (DFT) with the B3LYP functional and the 6-31G*+ basis set. For phosphorus-containing compounds (organophosphates), the LACVP*+ basis set was employed to account for effective core potentials. Frequency analyses were performed to confirm that all optimized geometries corresponded to true minima. Bond dissociation energies (BDEs) were computed as the enthalpy difference between the optimized parent molecule and the corresponding radical fragments. Specifically, BDE was calculated using the expression:

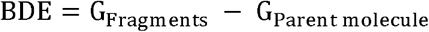

where G represents the Gibbs free energy.

### ML consensus rescoring method

The ML consensus rescoring method employed the Random□Forest [33] regression model implemented within the scikit-learn 1.6.1 Python package [51].□Each of the *T* decision trees in this model’s ensemble was independently trained to predict logIC_50_ based on the docking scores associated with the compound of interest, with mean squared error used as the trees’ training criterion. The final score output by the ensemble was obtained by taking the mean of its individual estimates. *T*, along with other hyperparameters associated with the ML consensus, was set via cross-validation for each docking score configuration (namely, the non-covalent and three covalent configurations).

*Training-testing dataset construction and weighting:* of the 412 inhibitors available, 226 (≈55□%) were allocated to training and the remaining 186 to testing, stratified by the 12 Murcko-based inhibitor clusters to preserve chemotype diversity. Because the experimental potencies were unevenly distributed, a kernel-density-estimation (KDE)-derived weighting scheme was applied during training to encourage the Random Forest to more equitably model structure-activity relationships across the complete observed range of biological response. Denote the docking score feature matrix by **x** and the associated logIC_50_ values by **y**. The Gaussian kernel-density estimator with bandwidth *h* yielded an estimated probability density 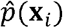 for each training sample *i*; its training weight was then

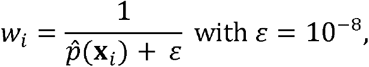

followed by normalization by the mean of all weights. This emphasized compounds residing in sparsely populated potency regions (e.g very strongly- and very weakly-/non-binding compounds).

*Hyperparameter optimization:* the ML consensus’ hyperparameters (Table 4) were optimized for each docking score configuration across 100 trials of Bayesian search, using the Tree-Parzen Estimator sampler from the Optuna 4.3.0 Python package [52]. Each trial was evaluated by five-fold cross-validation (CV), treating compound prioritization as a ranking problem and using Spearman’s r calculated over the predicted scores and experimental potencies as the optimization criterion:

**Table 4.** Hyperparameters associated with the ML consensus.

| Hyperparameter | Considered values | Remarks |
| --- | --- | --- |
| KDE bandwidth ($h$) | [0.1, 2.0] | Balances sensitivity to local variations (low values) and global trends (high values) in sample weighting. |
| Number of trees ($T$) | $[50, 300] \cap \mathbb{Z}$ | Fewer trees reduce computation, while more trees may improve stability and performance. |
| Maximum tree depth | $[2, 20] \cap \mathbb{Z}$ | Shallower trees may generalize better; deeper trees may capture more complex patterns. |
| Minimum number of samples required to split an internal node | $[2, 10] \cap \mathbb{Z}$ | Lower thresholds allow finer splits; higher thresholds promote simpler models. |
| Minimum number of samples required per leaf node | $[1, 5] \cap \mathbb{Z}$ | Smaller leaves capture more detail; larger leaves may yield more robust estimates. |

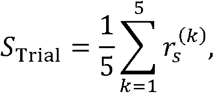

where 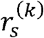 was the Spearman’s rank correlation on validation fold *k*.

*Final model fitting and prediction:* after the hyperparameter set optimizing cross-validated r under each docking score configuration was determined, the Random Forest was re-trained on the configuration’s complete training dataset with KDE weighting. The resulting model was used to generate consensus activity estimates for testing dataset compounds. Apart from comparisons to other methods in terms of r, the ML consensus was also considered in terms of its SHAP (SHapley Additive exPlanations) values to interrogate feature importance across structures, providing the consensus rankings with mechanistic interpretability. SHAP values decomposed each prediction into structure-specific contributions. Positive SHAP values indicated that a covariate increased the predicted potency for a given compound, while negative values reduced it, all relative to the model’s baseline. The magnitude of a structure’s SHAP values reflected its relative influence on the model’s output, enabling a visual readout of which structures most strongly guided compound prioritization under each docking score configuration. SHAP value analysis was conducted over the testing dataset using the TreeExplainer functionality [53] from the shap 0.46.0 Python package [54].

### CovDock score reweighting procedure

The CovDock score associated with a compound is the mean of its pre- and post-reaction complex docking scores. Since these scores may not be equally informative, a reweighted aggregation of both docking scores was devised. Let 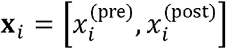 denote the two-dimensional vector of docking scores for compound *i*, where 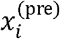 and 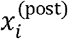 represent its pre- and post-reaction complex docking scores, respectively. The reweighted, aggregated docking score was defined as a linear combination of these components:

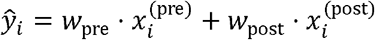

where the weights *w*_pre_ and *w*_post_ satisfy *w*_pre_ + *w*_post_ = 1. These weights were not preset but rather learned from the training dataset to maximize r between the reweighted scores and experimental potencies.

*Cross-validation and weight selection:* to determine optimal weightings, a grid search was conducted using a non-standard five-fold CV over all weight pairs within 𝒲, defined as:

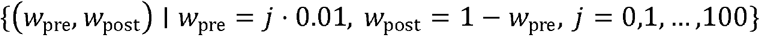

The weight pair yielding the greatest rank correlation 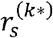 on the training (rather than validation) folds was selected as the representative weight pair of iteration *k* of the CV.

A single weight pair vector 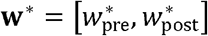 to use in the reweighting procedure was then created by taking the weighted mean of the five CV pairs:

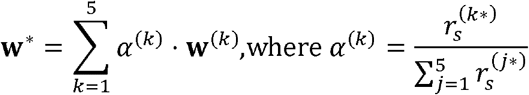

This weighting scheme employed between pairs emphasized those with stronger ranking performance over their respective CV iteration. The final weight pair vector w* was then used to compute reweighted aggregate scores on the held-out testing compounds:

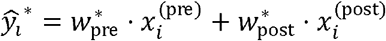

This method preserved interpretability while allowing the relative importance of pre- and post-reaction complex docking scores to be data-adaptive. The resulting reweighted aggregate docking scores were benchmarked against experimental potencies using r, consistent with the criteria used throughout this study.

## Author contributions

JCK: conceptualization, molecular docking and chemical analysis, visualization, writing – original draft. AAR: conceptualization, machine learning analysis, visualization, writing – original draft. YXH and HF: conceptualization, resources, writing – review & editing, supervision, project administration, funding acquisition.

## Data availability

The dataset utilized in this study, along with the code implementing the single- and multi-structure docking strategies, are available for access at https://github.com/bii-dpi/AChE_ML_consensus.

## Supplementary information

**Figure S1.**
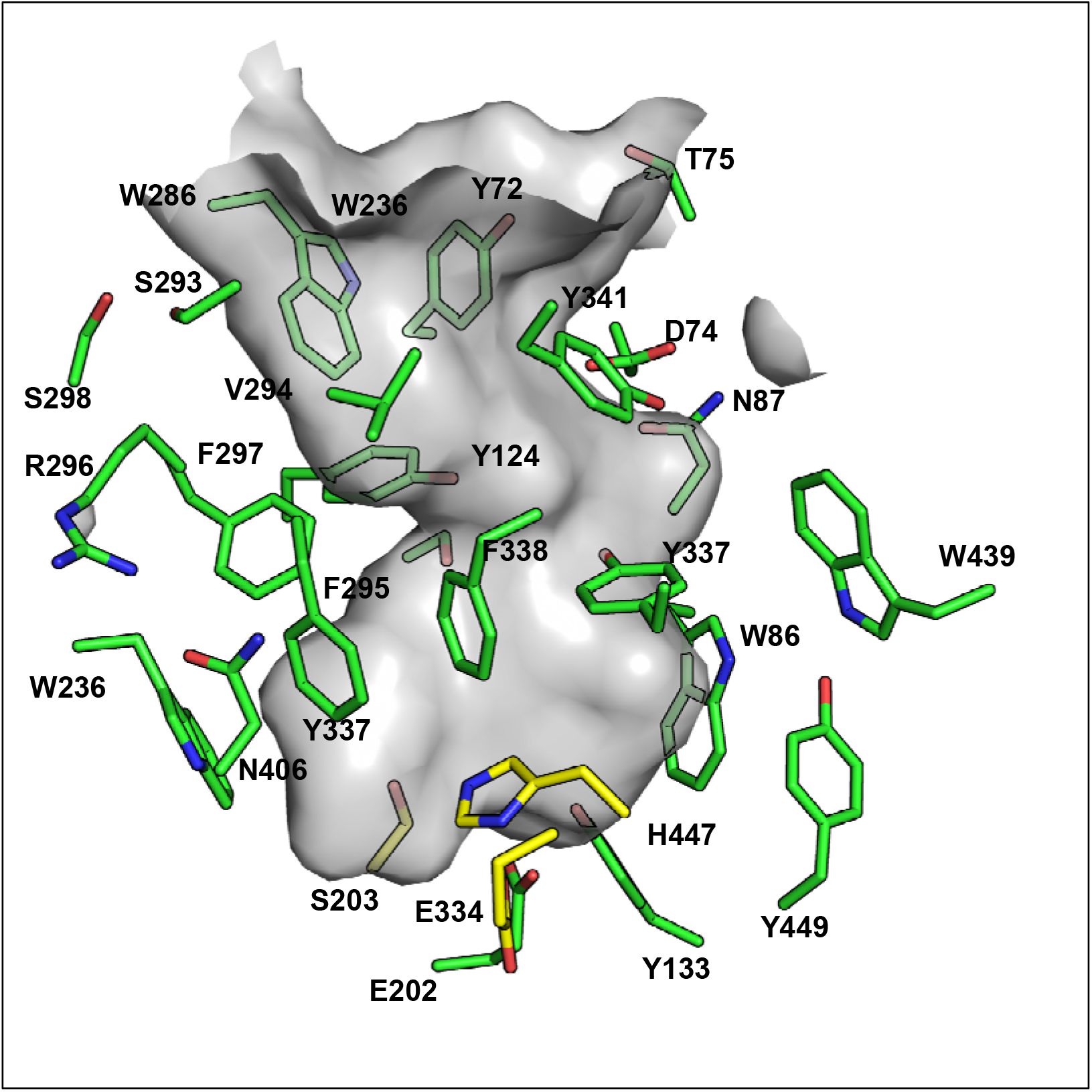
Binding-site residues of AChE within a 5 Å radius of the binding region, used for clustering.

**Figure S2.**
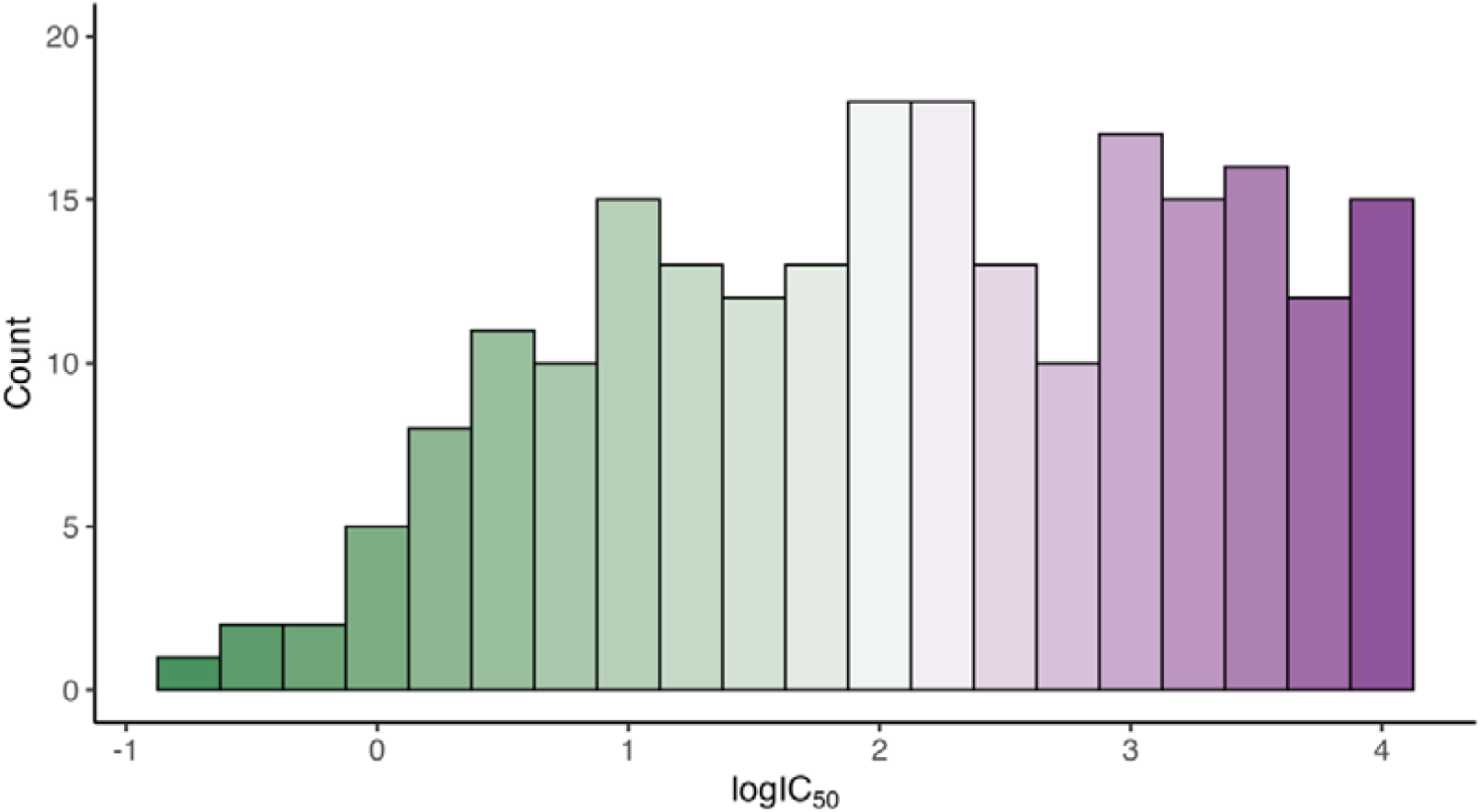
Histogram of potencies (logIC_50_) across the 226 compounds used for ML consensus training. Extreme potencies, particularly those of strong inhibitors, were underrepresented relative to compounds with moderate potency.

**Figure S3.**
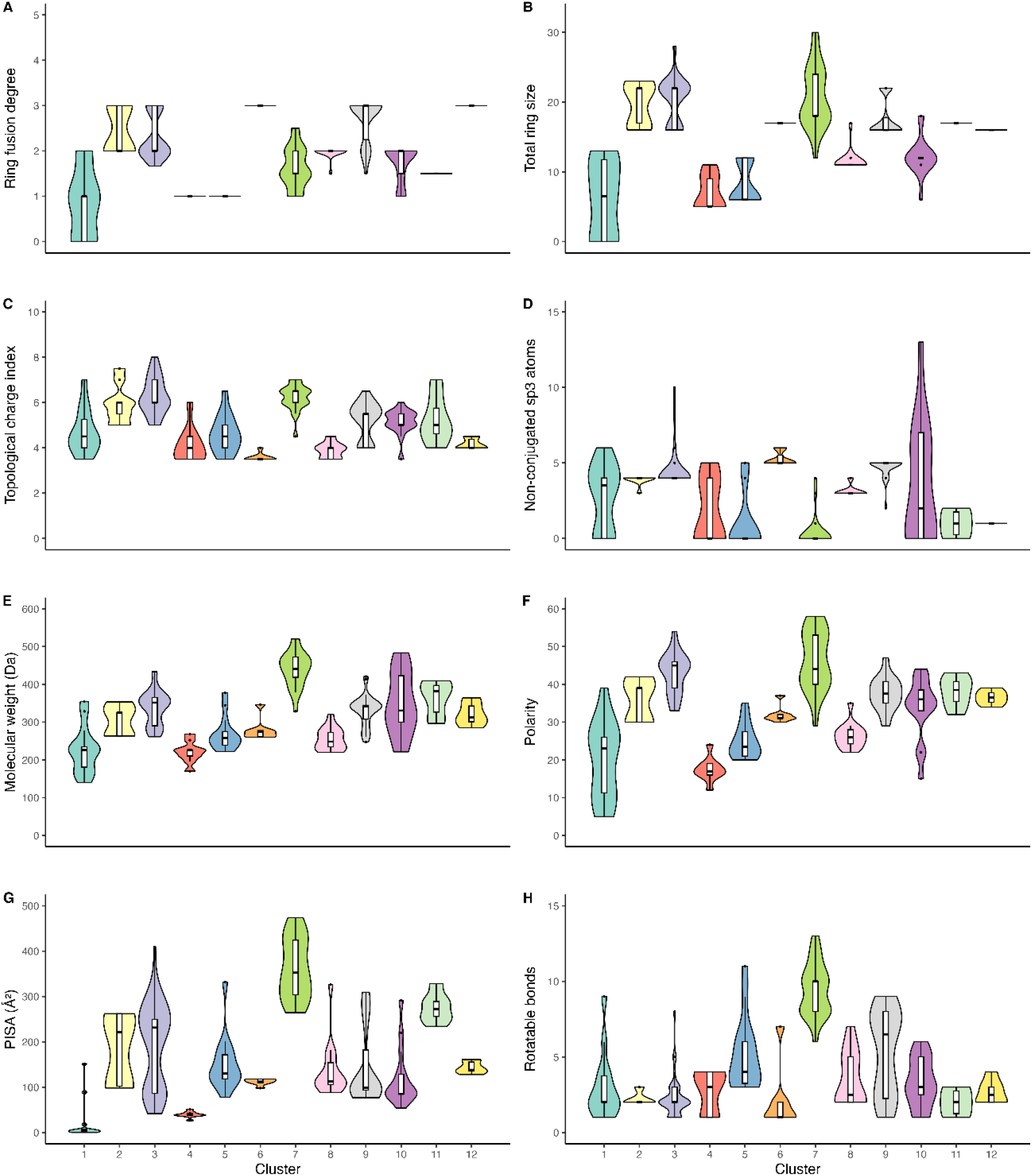
Distribution of ring fusion degree, total ring size, topological charge, nonconjugated C_sp3_ atoms molecular weight, polarity, π-surface area (PISA), and rotatable bonds across the chemical clusters.

**Table S1.** AChE inhibitor chemical cluster sizes.

| Cluster | Number of compounds |
| --- | --- |
| 1 | 30 |
| 2 | 21 |
| 3 | 90 |
| 4 | 20 |
| 5 | 32 |
| 6 | 15 |
| 7 | 46 |
| 8 | 31 |
| 9 | 66 |
| 10 | 33 |
| 11 | 14 |
| 12 | 14 |

**Table S2.** Performance comparison between CovDock scores and their reweighted alternative, as determined by cross-validation over the training dataset. Pre- and post-reaction weights may not add up to exactly one given their rounding.

| PDB ID | Pre-reaction<br>score weight | Post-reaction score<br>weight | Original CovDock<br>r | Rewighted<br>scoring r | Reweighting<br>improvement |
| --- | --- | --- | --- | --- | --- |
| 1B41 | 0.617 | 0.383 | 0.159 | 0.156 | -0.003 |
| 4M0E | 0.348 | 0.652 | 0.187 | 0.183 | -0.004 |
| 6NTO | 0.360 | 0.640 | 0.262 | 0.262 | -0.001 |
| 6WUY | 0.310 | 0.690 | 0.534 | 0.551 | 0.017 |
| 8AEN | 0.342 | 0.658 | 0.434 | 0.430 | -0.004 |

**Table S3.**
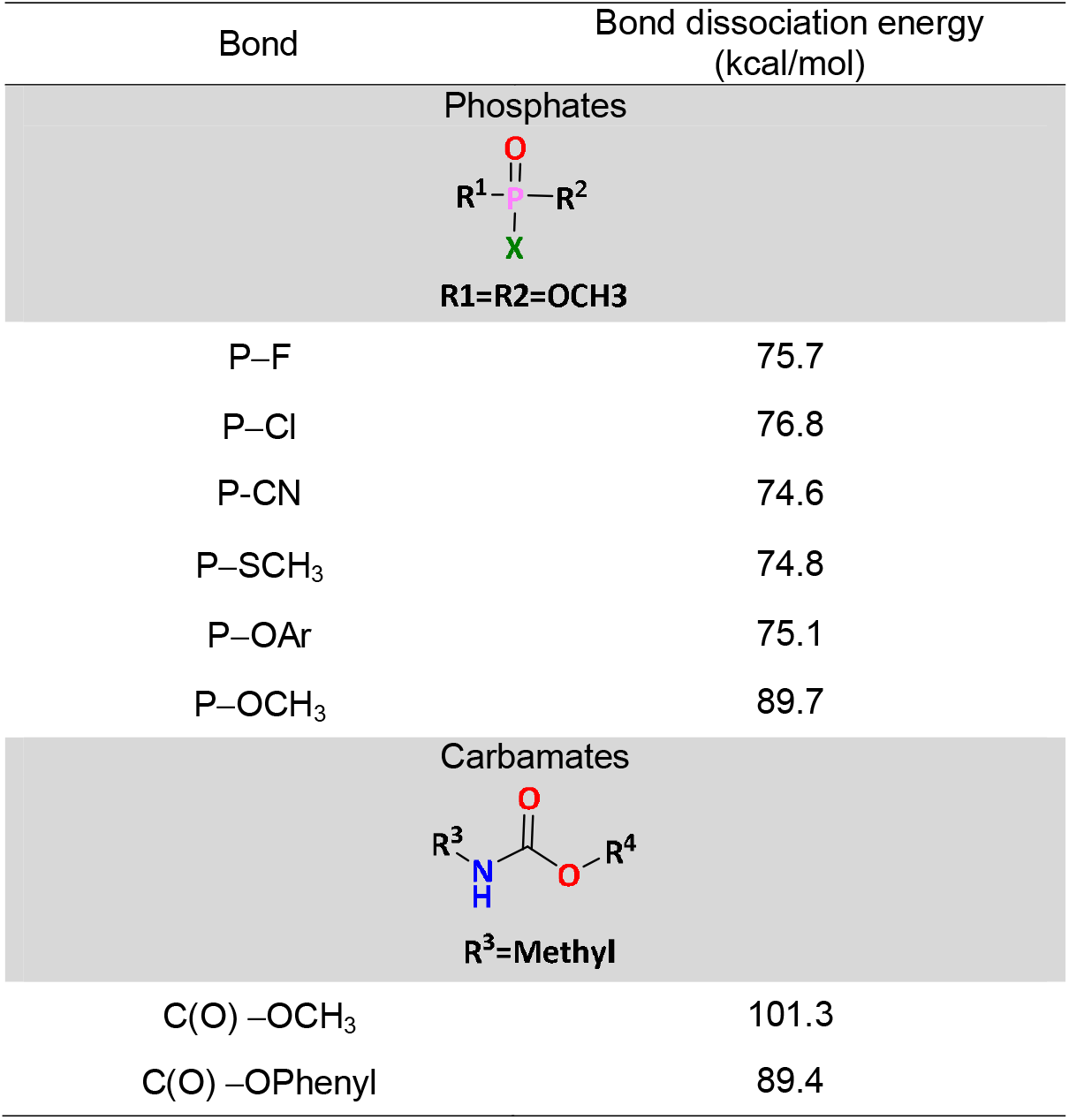
Bond dissociation energies of organophosphate bonds using density functional theory with basis set LACVP 6-31 G+*.

